# Cell proliferation and Notch signaling coordinate the formation of epithelial folds in the *Drosophila* leg

**DOI:** 10.1101/2023.09.28.559944

**Authors:** Alonso Rodríguez, Sergio Córdoba, Daniel Felipe-Cordero, Antonio Baonza, David Miguez, Carlos Estella

## Abstract

The formation of complex three-dimensional organs during development requires the precise coordination between patterning networks and mechanical forces. In particular, tissue folding is a crucial process that relies on a combination of local and tissue-wide mechanical forces. Here, we investigate the contribution of cell proliferation to epithelial morphogenesis using the *Drosophila* leg tarsal folds as a model. We reveal that tissue-wide compression forces generated by cell proliferation, in coordination with Notch signaling pathway, are essential for the formation of epithelial folds in precise locations along the proximo-distal axis of the leg. As cell numbers increase, compressive stresses arise, promoting the buckling of the epithelium and reinforcing the apical constriction of invaginating cells. Additionally, the Notch target *dysfusion* (*dysf*) plays a key function specifying the location of the folds, through the apical accumulation of F-actin and the apico-basal shortening of invaginating cells. These findings provide new insights into the intricate mechanisms involved in epithelial morphogenesis, highlighting the critical role of tissue-wide forces in shaping a three-dimensional organ in a reproducible manner.

## Introduction

The transition during development from a relatively flat epithelium to a complex three-dimensional structure such as the gut or the neural tube relies largely on tissue folding during morphogenesis. The precise location of the folds that sculpt the shape of an organ is dictated by the underlying patterning network, and changes in cell shape and mechanical forces, such as tension and compression, drive the process (1). Asymmetrical mechanical forces generated at both local and tissue-wide levels are the main inductors of the cell shape changes that promote epithelial folding (2). Many different mechanisms, including apical constriction (3), differential planar cell proliferation (4), apoptotic basal extrusion (5), mitotic cell rounding (6) or basal relaxation and lateral tension (7) contribute to the generation of these local forces. In addition, differential growth between tissue layers or within the plane of the tissue may also contribute to tissue folding through buckling (4) (8, 9) (reviewed in (2)). An important question is how these local and tissue-wide mechanisms are coordinated to drive epithelial folding in the highly reproducible pattern that shape developing tissues and organs.

The imaginal discs of *Drosophila* are an excellent model to study morphogenesis as the epithelium folds in a stereotyped and reproducible manner, and their underlying patterning mechanisms are extensively studied (10). The imaginal discs are epithelial sac-like structures formed by a monolayer of pseudostratified cells that during larval development grow and become progressively subdivided in territories with different cellular identities. At metamorphosis, imaginal discs give rise to adult external structures such as the wing or the leg (10). The leg imaginal disc is divided into ten segments along the proximo-distal axis, each segment being determined by the expression of a distinct code of transcription factors that confer identity to each segment and dictate the positioning of the Notch ligands Delta (Dl) and Serrate (Ser) (10-12). The activation of Notch in nine rings at the distal end of each leg segment not only directs the formation of the epithelial folds that prefigure the adult joints but also contributes no-autonomously to the growth of the appendage (12-15). At the tarsal segments, the Notch target gene *dysfusion* (*dysf*) controls epithelial fold morphogenesis and adult joint formation through the spatial regulation of Rho1 activity (16, 17). The Rho1 GTPase induces acto-myosin contractility at the cell apex. The anchor of F-actin at the adherens junctions transmits the contractility to neighboring cells, resulting in tissue invagination (3). In addition, it has been proposed that apico-basal forces generated by patterned apoptotic cells could drive epithelium folding (18, 19). All these studies describe the molecular and cellular mechanisms implicated in the formation of the epithelial fold morphogenesis at the local scale, that is at the cells that invaginate. However, the impact that tissue-wide forces such as those generated by cell proliferation have on shaping these folds remains mostly unexplored.

The aim of this study is to investigate the contribution of cell division to epithelial morphogenesis, using *Drosophila* leg disc tarsal folds as a model. We have found that cell proliferation generates tissue-wide compression forces that, in coordination with the Notch pathway, drives the formation of the epithelial folds. Our findings indicate that the Notch target *dysf* plays a crucial role in specifying the location of the tarsal epithelial folds. This involves providing cells with unique mechanical properties that lead to the apical accumulation of F-actin and the apico-basal shortening of invaginating cells. Additionally, we describe that cell proliferation generates compressive stresses, which contribute to the buckling of the epithelium through the apico-basal elongation of interjoint cells, while simultaneously promoting the apical constriction of invaginating cells. In addition, we used a simple computer-based simulation model that recreated the initial process of tarsal fold formation. This model accurately predicts the folding phenotypes and the behavior of cells, even when cell proliferation is reduced or when the Dysf factor is absent.

Overall, our results provide new insights into the intricate mechanisms underlying epithelial morphogenesis where multiple mechanisms act together to sculpt a three-dimensional organ in a precise and reproducible manner.

## Results

### Dynamics of leg epithelial fold formation

To study the contribution of cell proliferation during epithelial folding we use the *Drosophila* leg imaginal disc as a model. This disc is a sac-like structure composed of two different cell types, a monolayer of pseudostratified epithelial columnar cells that will give rise to the adult appendage and a monolayer of squamous cells that confine the main epithelium, named peripodial membrane. In addition, an extracellular matrix (ECM) provides an elastic constraining environment that contribute to shape the appendages (20, 21). The imaginal epithelium of the third instar larval leg disc is characterized by concentric folding along the proximo-distal (PD) axis, which prefigures the division of the adult leg into ten distinct segments that are separated by flexible joints (12, 14). To explore the relationship between cell proliferation and epithelial folding during development, we used phalloidin staining to visualize F-actin distribution in the folding cells, as well as the mitotic marker phospho-histone 3 (pH3) and EdU (5-ethynyl-2’-deoxyuridine) labeling to identify actively dividing cells. We also stained imaginal discs with a *lacZ* reporter for the tarsal Notch target gene *dysf* (15, 17). From mid-third instar larval stage to prepupa, we observed a sequential accumulation of F-actin in ring-like structures at the folding points, which ultimately form the joints separating the leg segments in the adult stage (Fig. 1A). This stereotyped pattern of epithelial folding follows the reported sequential and reproducible activation of the Notch pathway during imaginal disc development (12, 14, 22). Dividing cells are found throughout the leg imaginal discs with no particular cell division pattern and their number progressively decreases during early prepupal stages (Fig. 1A and S1 Fig). The distal folds that prefigure the formation of the adult tarsal joints initiate at mid-third instar and are fully recognizable in later stages by F-actin accumulation distal to *dysf* expression (Fig. 1B and C). It is important to highlight that while the apical side of the leg epithelium is deformed during the formation of the tarsal folds, the basal side maintains a remarkably straight line as cells are strongly attached to the basal matrix (Fig. 1C and see below).

**Figure 1:**
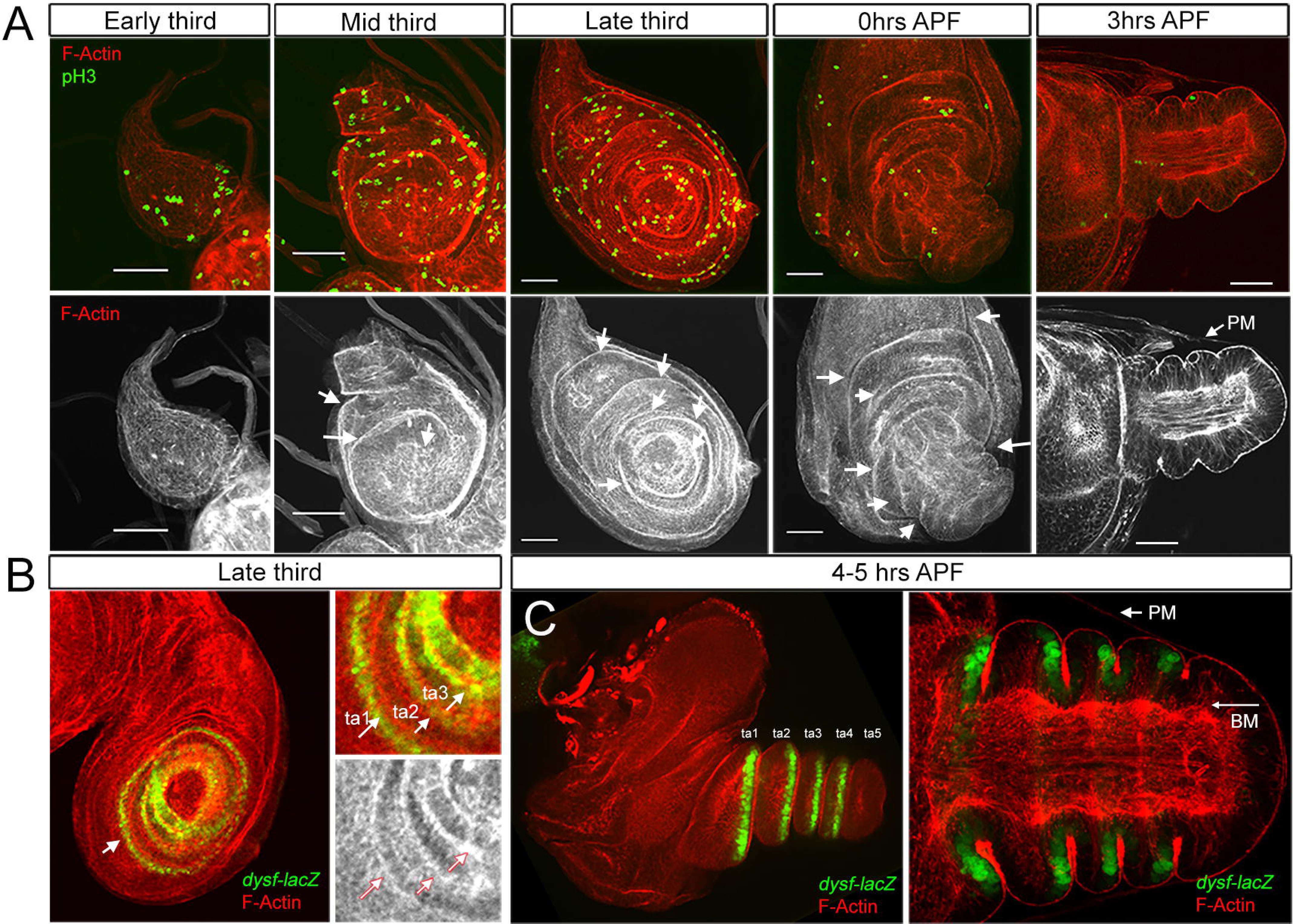
Dynamics of cell proliferation and fold formation in the leg. A) Time course of the leg imaginal disc at different time points of development and stained for Phalloidin (F-actin, red and gray) and pH3 (green). Single channel for F-actin is also shown. Arrows indicate the formation of the epithelial folds by F-actin accumulation and the peripodial membrane (pm) in the early pupa. Scale bars: 50 µm. B) Third instar leg imaginal discs stained for F-actin (red and white) and *dysf-lacZ* (green). In the right panels, the formation of some tarsal folds (ta1-ta3) is indicated by arrows. C) A 4-5 hrs APF leg imaginal disc stained for F-actin (red) and *dysf-lacZ* (green). In the right panel a close-up view of the tarsal segments is shown in a sagittal section. The arrow indicates the basal membrane (BM) and the peripodial membrane (PM).

### Cell proliferation is required for leg epithelial folding and adult joint formation

As leg epithelial folding is taking place in a highly proliferative tissue, we decided to investigate the possible influence of cell division on tarsal epithelial morphogenesis. To this end, we employed a combination of the *hedgehog* (*hh*)*-Gal4* driver and the *tub-Gal80^ts^* (*hh^Gal80^>*) and *UAS-E2f1* RNAi constructs to specifically and temporally knock down the expression of E2f1, a key regulator of the cell cycle (23), in the posterior compartment at defined time-points. This was achieved through the expression of a specific RNAi line, and the knockdown was conducted for a duration of 48 hours (hrs) during mid-third instar larval development (Fig 2A and S2A-B Fig). We found that this treatment reduces about 35% the number of cells in the posterior distal leg domain when compared to the anterior control compartment. For this, we measured the number of nuclei at the tarsal domain in several planes in the posterior and anterior compartments. However, we did not observe a reduction in the mitotic index in the wing disc, calculated as the number of pH3-positive cells per area, suggesting that cells are proliferating at a slower rate (S2 Fig). The downregulation of E2f1 during mid-third instar larval development has a strong effect on the formation of the tarsal epithelial folds, more apparent at the prepupal stage (Fig. 2A and B): while in the control anterior compartment the four characteristic tarsal folds are observed, in the posterior compartment the epithelium flattens and adult tarsal joints are lost (Fig. 2A and B). To confirm our findings, we suppress E2f1 function using two additional approaches: by expressing an activated form of the E2f1 repressor Retinoblastoma (Rbf^280^) (24) and by knocking down the E2f1 target gene CycE (23). Consistent with our previous observations, both methods produced comparable results (Figure 2A and B).

**Figure 2:**
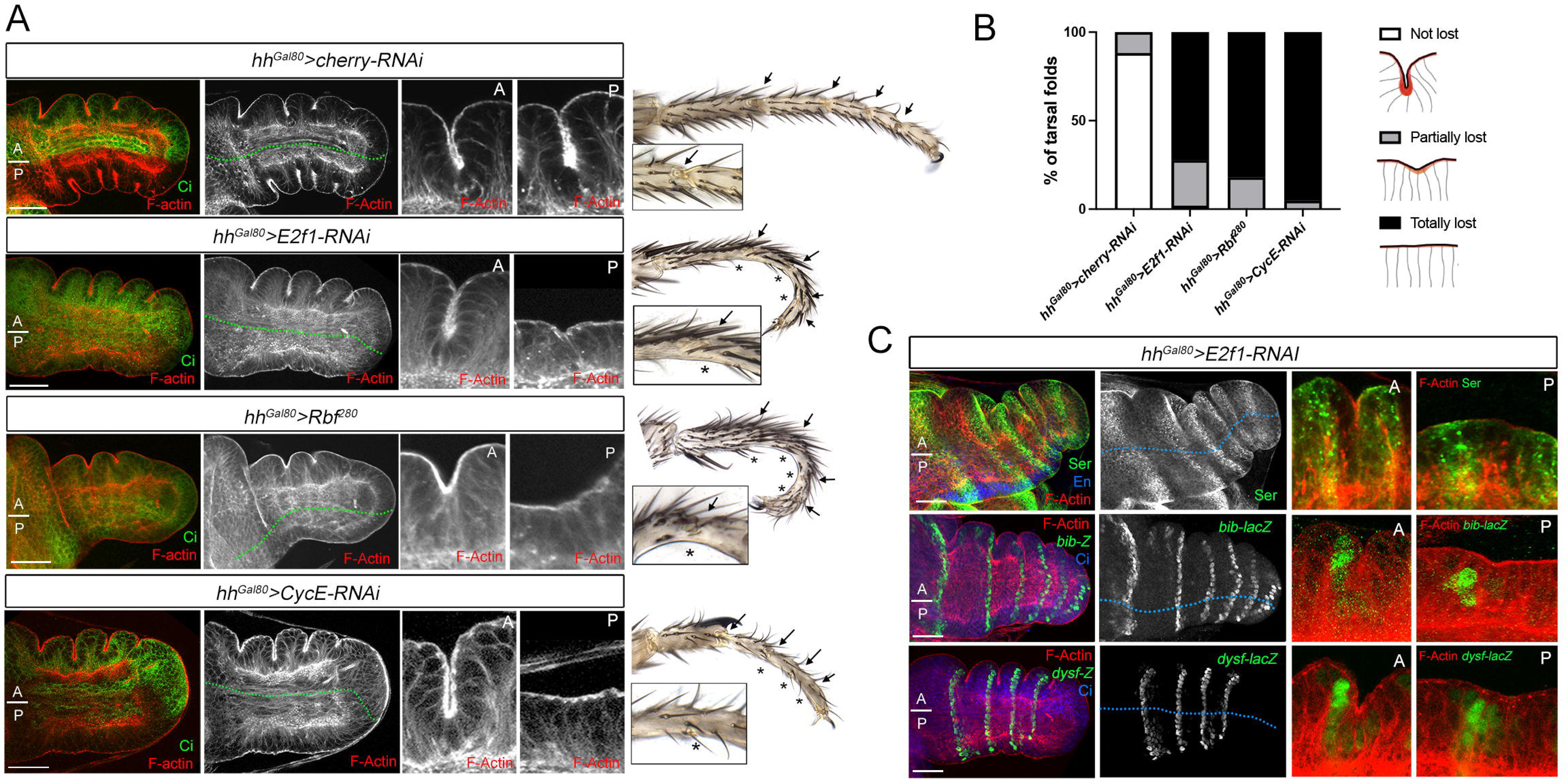
The reduction of cell proliferation inhibits epithelial folding and joint formation. A) Tarsal region of prepupal leg imaginal discs expressing the indicated transgenes for 48 hrs by the *hh-Gal4*, *tub-Gal80^ts^* (*hh^Gal80^>*) stained for F-actin (red and white), and Cubitus interruptus (Ci, green) that marks the anterior compartment. The *cherry-RNAi* line was used as a control. The anterior and posterior compartments are indicated as A and P, respectively and their boundary is represented by a dotted green line. A higher magnification of an anterior and posterior sagittal view of tarsal folds is presented for each genotype. The respective adult legs phenotypes of these experiments are shown with the tarsal joints indicated as arrows and the affected ones by asterisks. A representative close view of a tarsal joint is also presented. B) Quantification of tarsal fold defects for the prepupal legs shown in A. Severity of the phenotypes is assessed by counting the number of tarsal joints that presented folding defects: *‘not lost’* if the tarsal folds are not affected; *‘partially lost’* when the tarsal fold is recognized but reduced compared to the respective control one; ‘*lost’* when the tarsal fold is not present. The genotypes are as follows: control *hh^Gal80^>cherry-RNAi* (24 legs, 91 folds)*, hh^Gal80^>E2f1-RNAi* (37 legs, 101 folds), *hh^Gal80^>Rbf^280^* (21 legs, 61 folds), *hh^Gal80^>CycE-RNAi* (7 legs, 21 folds). The larvae from these genotypes were kept at 17°C and transferred to 31°C at the beginning of third instar. C) Prepupal leg imaginal discs of the *hh^Gal80^>E2f1-RNAi* genotype dissected 48 hrs after inducing the transgene in the posterior compartment and stained for F-actin, Ser (green), *bib-lacZ* (green) and *dysf-lacZ* (green). Engrailed (En, blue) and Ci (blue) staining are used to indicate the posterior (P) and anterior (A) compartments, respectively. The antero-posterior compartment boundary is represented by a dotted blue line. A higher magnification of an anterior and posterior tarsal fold is indicated for each staining. Scale bars: 50 µm.

The absence of leg epithelial folding observed upon slowing down cell proliferation may be attributed to a failure in joint specification. To test this possibility, we stained for the Notch ligand Ser and for two Notch target genes: *big brain* (*bib*) and *dysf* (12, 14, 17). While the knockdown of E2f1 in the posterior compartment prevents the folding of the epithelium, no defects in the expression of *Ser*, *bib* or *dysf* are observed (Fig. 2C). These results indicate that the absence of leg folds observed after reducing cell proliferation is not a direct consequence of the lack of Notch activity.

To extend these results we used the *apterous* (*ap*) Gal4 line (*ap^42B11^*) to reduce cell division in a restricted tarsal domain of the leg, the fourth tarsal segment encompassing the t4/t5 fold (S3 Fig). The knockdown of E2f1 or CycE, or the expression of *Rbf^280^* in this domain has a strong effect on the formation of the adult t4/t5 joint without affecting *dysf-lacZ* expression, a Notch activity reporter (17) (S3 Fig).

To investigate whether cell division is specifically required in the cells involved in the folding process or if overall tissue-level cell proliferation contributes to epithelial folding, we utilized the *dysf-Gal4* line to knockdown E2f1. This line is strongly activated in the cells that will form the tarsal folds, starting at third instar larval stage (17). Tarsal epithelial folds and adult joints were still formed in these experimental conditions (S4 Fig). We observed the lack of some bristles in the tarsal segments due to the late residual activity of the *dysf-Gal4* line in the interjoint domain and the requirement of cell division in the sensory organ precursors (25). Similar results were obtained when the *bib-Gal4* line, which also drives expression specifically in folding cells (26), was used to reduce cell proliferation (S4 Fig).

Taken together, our results suggest that cell proliferation is required for epithelial folding and adult tarsal joint formation. Importantly, this requirement is not specifically needed at the folding cells.

### Rho1 activity and apical constriction at the tarsal epithelial folds are impaired in proliferation deficient cells

Localized activation of Rho1 by Dysf in the tarsal segments promotes epithelial folding through cell shape changes such as apical constriction (16). Apical constriction is mediated by the shrinkage of the apical cortex surface by the actin-myosin networks and the generated forces are transmitted to the neighboring cells through the adherens junctions’ attachments (3). To investigate the impact of cell proliferation on the cell shape changes necessary for epithelial folding, we downregulated E2f1 specifically in the posterior compartment (using *hh^Gal80^>E2f1-RNAi*) and examined apical constriction through E-Cadherin and F-actin staining. We observed that posterior cells at the presumptive joint fold failed to properly constrict apically and show significantly less apical F-actin than control anterior cells (Fig. 3A, B and E). Next, we monitored Rho1 activity using a GFP-based sensor (Rho1-BD-GFP) which contains the three Rho1 binding domains of Protein kinase N fused to GFP and that recognizes the active form of Rho1 (27). Expression of the *UAS-Rho1-BD-GFP* construct under the *hh-Gal4* driver shows a strong accumulation of GFP in the tarsal folds and specifically at the apical domain of the cells that apically constrict (Fig. 3C)(16). Importantly, GFP accumulation, and therefore Rho1 activity, is reduced but not eliminated in posterior tarsal cells of the fold region where cell proliferation was decreased by the knockdown of E2f1 (Fig. 3D and F). Next, we followed Myosin II regulatory light chain localization using a GFP-tagged version of *spaghetti squash* (Sqh-GFP). As previously described (16), Sqh is located subapically at the level of the adherens junctions and although the apical distribution does not change after E2f1 knockdown, we observed less accumulation than in control anterior cells (Fig. 3G).

**Figure 3:**
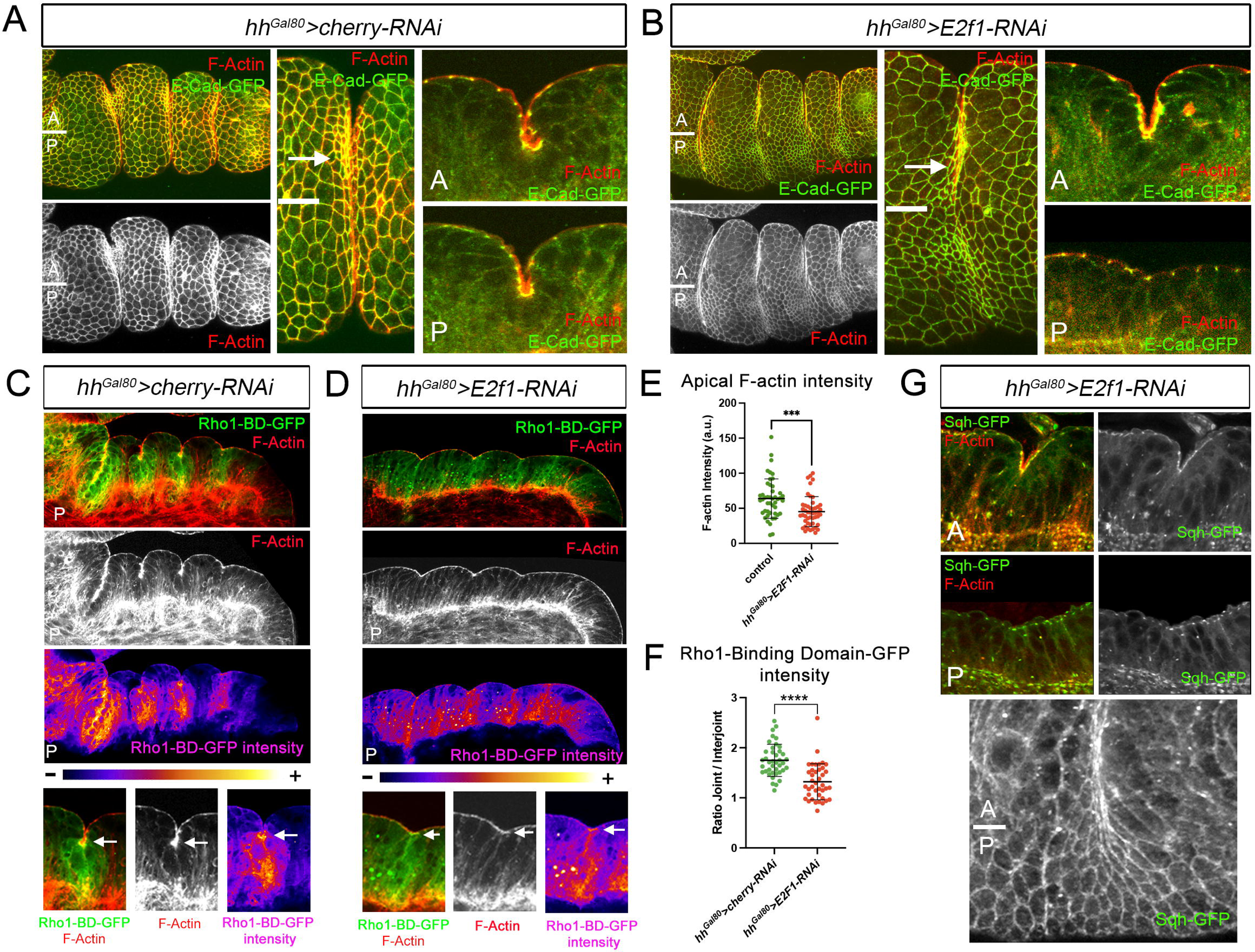
Analysis of F-actin, Rho1 activity and Sqh after E2f1 knock down. A-B) Tarsal region of prepupal leg imaginal discs expressing for 48 hrs the indicated transgenes by the *hh-Gal4*, *tub-Gal80^ts^* (*hh^Gal80^>*) stained for F-actin (red and gray), and E-Cad (green). The anterior and posterior compartments are indicated as A and P, respectively and their boundary is represented by white line. A higher magnification of an anterior and posterior sagittal view of tarsal folds is presented for each genotype. An arrow indicates the accumulation of F-actin in the control anterior compartment cells that are apically constricted. C-D) Posterior tarsal region of prepupal leg imaginal discs expressing for 48 hrs the indicated transgenes by the *hh-Gal4*, *tub-Gal80^ts^* (*hh^Gal80^>*) and the *Rho1-BD-GFP* (green and magenta) stained for F-actin (red and white). Regions of enhanced GFP levels are seen around the joints in C that are separated by regions of lower GFP levels in the interjoint regions. This pattern is lost in *hh^Gal80^>E2f1-RNAi*, where GFP levels accumulation in the folding region is reduced. Close up views of a sagittal section a posterior tarsal fold of the genotypes indicated above showing separate channel for F-actin (gray) and false color to enhance contrast to the right. GFP and F-actin levels are accumulated apically in C in the cells that are undergoing apical constriction (arrow), while in D, less accumulation of GFP and F-actin is observed. E) Quantification of apical F-actin (mean intensity) at the joint cells in *hh^Gal80^>cherry-RNAi* (control) and *hh^Gal80^>E2f1-RNAi* prepupal legs. A total of 16 legs and 46 joints were quantified per genotype. ***p<0.0005, with Student’s t test, indicating a significant difference from control. Error bars represent standard deviation (SD). F) Ratio of fluorescence levels of Rho1RBD-GFP (mean intensity) within joint and interjoint domains in *hh^Gal80^>cherry-RNAi* (control, 13 legs, 40 joints and interjoints) and *hh^Gal80^>E2f1-RNAi* (14 legs, 41 joints and interjoints) prepupal legs. A ratio of 1 would imply the same levels in joint and interjoint domains, while any increment over 1 means higher levels in the joint vs interjoint domain. ****p<0.0001, with Student’s t test, indicating a significant difference from control. Error bars represent SD. G) Higher magnification of an anterior (A) and posterior (P) sagittal view of tarsal folds from *hh^Gal80^>E2f1-RNAi* prepupal legs stained for F-actin (red and white) and Sqh-GFP (green and white). An apical view of a tarsal joint where the anterior and posterior compartments are indicated showing the accumulation of Sqh-GFP (white) in the control anterior compartment cells that are apically constricted.

These results suggest that cell proliferation, together with Dysf, contributes to Rho1 activation, that in turn promotes the actin-myosin network contraction and apical constriction of the cells that form the presumptive joint fold.

### Tissue compression generated by cell proliferation is necessary for leg epithelial folding

Our previous results indicate that cell division contributes to the folding of the leg imaginal epithelium through Rho1 mediated apical constriction at specific locations dictated by Dysf. This suggests that compressive stresses generated during cell division in the imaginal disc could contribute to folding the epithelium by buckling. To gain a more comprehensive understanding of the folding mechanism in the leg epithelium, we measured the heights of cells in the interjoint and joint domains before and after the formation of the t4/t5 fold. Specifically, we conducted these measurements over a time frame of 0 to 6 hours after puparium formation (APF). Although this particular tarsal fold is the last one to be shaped, our results indicate that the leg imaginal epithelium also requires cell division to properly fold and to form the corresponding adult joint (S3 Fig). While the height of the cells that form the joint fold (distal to *dysf* positive cells) gradually decreases as the epithelium invaginates (Fig. 4A and B), we observed that the interjoint cells (proximal to *dysf* positive cells) progressively increased their heights until approximately five hours APF when it suddenly drops (Fig. 4A and B). This particular time point (6 hrs APF) is concomitant to basal extracellular matrix degradation and the removal of the peripodial membrane that allows leg expansion (20, 28). The cellular height increase at the interjoint domain is appreciated by the long actin filaments that attach the cells to the basement membrane (Fig. 4A and C). These results indicate that the formation of tarsal folds involves not only a decrease in the height of the invaginating cells, but also a notable increase in the height of the cells located in the interjoint domain.

**Figure 4:**
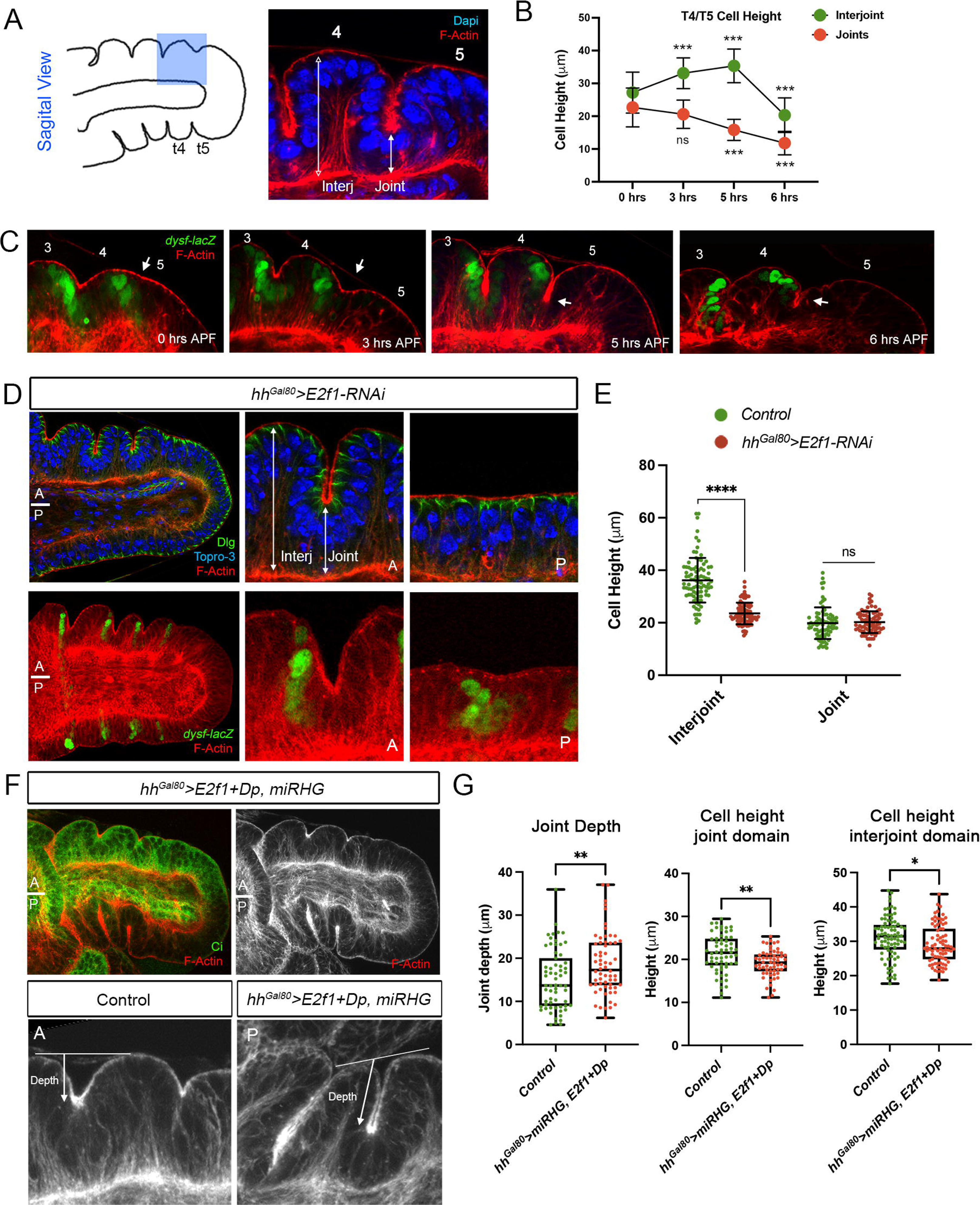
Cell proliferation promotes the buckling of the leg epithelium at specific locations. A) Schematic representation of a sagittal view of the tarsal domain of a prepupal leg with the region corresponding to the t4/t5 tarsal fold highlighted in blue. Staining of the t4/t5 tarsal fold with Dapi (blue) and F-actin (red). The interjoint and joint domains are indicated. B) Quantification of the cell heights of the t4/t5 interjoint (green) and joint (red) cells form 0 hrs to 6 hrs APF. 0 hrs: 34 joints, 34 interjoints. 3 hrs: 24 joints, 24 interjoints. 5 hrs: 26 joints, 26 interjoins. 6 hrs: 30 joints, 30 interjoints. Error bars represent standard deviation (SD). Statistical analysis by one-way ANOVA when compared the mean of each time point with the mean of the control (0 hrs) as indicated. ***<p=0.001 and ns=not significant. C) Time course imaging of the sagittal view of a t4/t5 joint (arrow) at different time points stained for F-actin (red) and *dysf-lacZ* (green). D) Prepupal leg imaginal discs of the *hh^Gal80^>E2f1-RNAi* genotype dissected 48 hrs after inducing the transgene in the posterior compartment and stained for F-actin (red), Dlg (green) or *dysf-lacZ* (green) and Topro-3 (blue). The antero-posterior compartment boundary is represented by a white line. A higher magnification of an anterior (A) and posterior (P) tarsal folds is indicated for each staining. E) Quantification of the cell heights at the interjoint and joint domains in *hh^Gal80^>cherry-RNAi* (control, 21 legs, 63 joints, 79 interjoints) and *hh^Gal80^>E2f1-RNAi* (21 legs, 62 joints, 78 interjoins) prepupal legs. ****p<0.0001 and ns=not significant, with Student’s t test, indicating a significant difference from control. Error bars represent SD. F) Prepupal leg imaginal discs of the *hh^Gal80^>E2f1+Dp, miRHG* genotype dissected 24-30 hrs after inducing the transgene in the posterior compartment and stained for F-actin (red and gray) and Ci (green). The antero-posterior compartment boundary is represented by a white line. A higher magnification of an anterior (A) and posterior (P) tarsal folds is indicated. The anterior compartment is used as a control. Also indicated is how the joints depth is measured. G) Quantification of joint depth and the cell heights at the interjoint and joint domains of the prepupal legs in F. A total of 22 legs were dissected and 65 joints and 87 interjoints were measured. **p<0.01, *p<0.05 and ns=not significant, with Student’s t test, indicating a significant difference from control. Error bars represent the minimum and maximum points.

Next, we analyzed the height of interjoint and joint cells after reducing cell proliferation by expressing the *E2f1-RNAi* line in the posterior compartment, and compared it with the anterior control in 3 hrs APF leg discs. As in the experiments above, we used *dysf-lacZ* as a marker for the region where the joints will form. We found that the knockdown of E2f1 led to a significant reduction in the height of interjoint cells. This, in turn, resulted in the suppression of the upward fold, as depicted in Fig. 4D and E. In contrast, the relative height of the joint cells does not significantly change when compared to the anterior control cells (Fig. 4D and E). These results suggest that compressive forces generated by cell proliferation promote the upward buckling of the interjoint epithelium and thus contribute to the formation of the tarsal folds.

Next, we analyzed the effects of an increase in cell proliferation on tissue folding. To this end, we ectopically expressed E2f1, and its obligated partner Dp, in the posterior compartment. As E2f1 not only promotes cell proliferation but also cell death, we inhibited apoptosis using the UAS-miRHG construct that efficiently represses the activity of the proapoptotic genes (29, 30). We also stimulated proliferation by the induction of the transcriptional effector of the Hippo pathway, Yorkie (Yki) (31). We employed the *hh^Gal80^* system to restrict the expression of these transgenes to 24-30 hours before pupae dissection (3 hrs APF). This was done to avoid deformation of the leg epithelium, as we have observed that prolonged expression times can prevent an accurate phenotypic analysis. In both experiments, although the overall morphology and location of the leg epithelial folds are maintained, we detected an increase on the depth of the invaginated joint region that is accompanied with a decrease on the heights of the cells in this domain when compared to the control anterior compartment (Fig. 4F-G and S5 Fig). However, contrary to what would be expected, we did not observe an expansion on the apico-basal height of the interjoint cells, suggesting that most probably these cells have reached the maximum on its elongation (Fig. 4E-G and S5 Fig).

These observations suggest that an increase in tissue compression resulting from cell proliferation increase epithelial folding at specific positions. In the case of the leg discs, these positions are dictated by Dysf.

### The extracellular matrix contributes to generate the compression forces that promote the folding of the epithelium

Imaginal discs grow surrounded by the ECM that act as elastic constraining element that helps shape them during morphogenesis (21)(20, 32). To investigate the role that the ECM has in promoting leg fold and joint formation we first analyzed Collagen-IV, a main component of the basal matrix, using a GFP-tagged version of the protein Viking (Vkg-GFP). As previously reported (20), we found that around 4 hrs APF, main epithelium cells are attached to the basal ECM (Fig. 5A). At later stages of pupal development (around 7 hrs APF), ECM remodeling and peripodial membrane removal initiate the elongation of the leg along the proximo-distal axis (Fig. 5A) (20, 21, 28, 33). Next, we tested whether removal of the basal ECM in the leg affects fold and adult joint formation. To this end we overexpressed the matrix metalloproteinase (MMP2) in the distal domain of the leg with the *rotund* (*rn*) Gal4 line (Fig. 5B). We found that the leg epithelial folds are highly disrupted as the distal appendage is prematurely expanded along the proximo-distal axis (Fig. 5B). Quantification of cell heights demonstrated the apico-basal shortening of interjoint cells in this experimental condition (Fig. 5C). Remarkably, basal ECM degradation also affects the formation of some joints in the adult leg (Fig. 5D). These results confirm the role of the ECM promoting epithelial folding.

**Figure 5:**
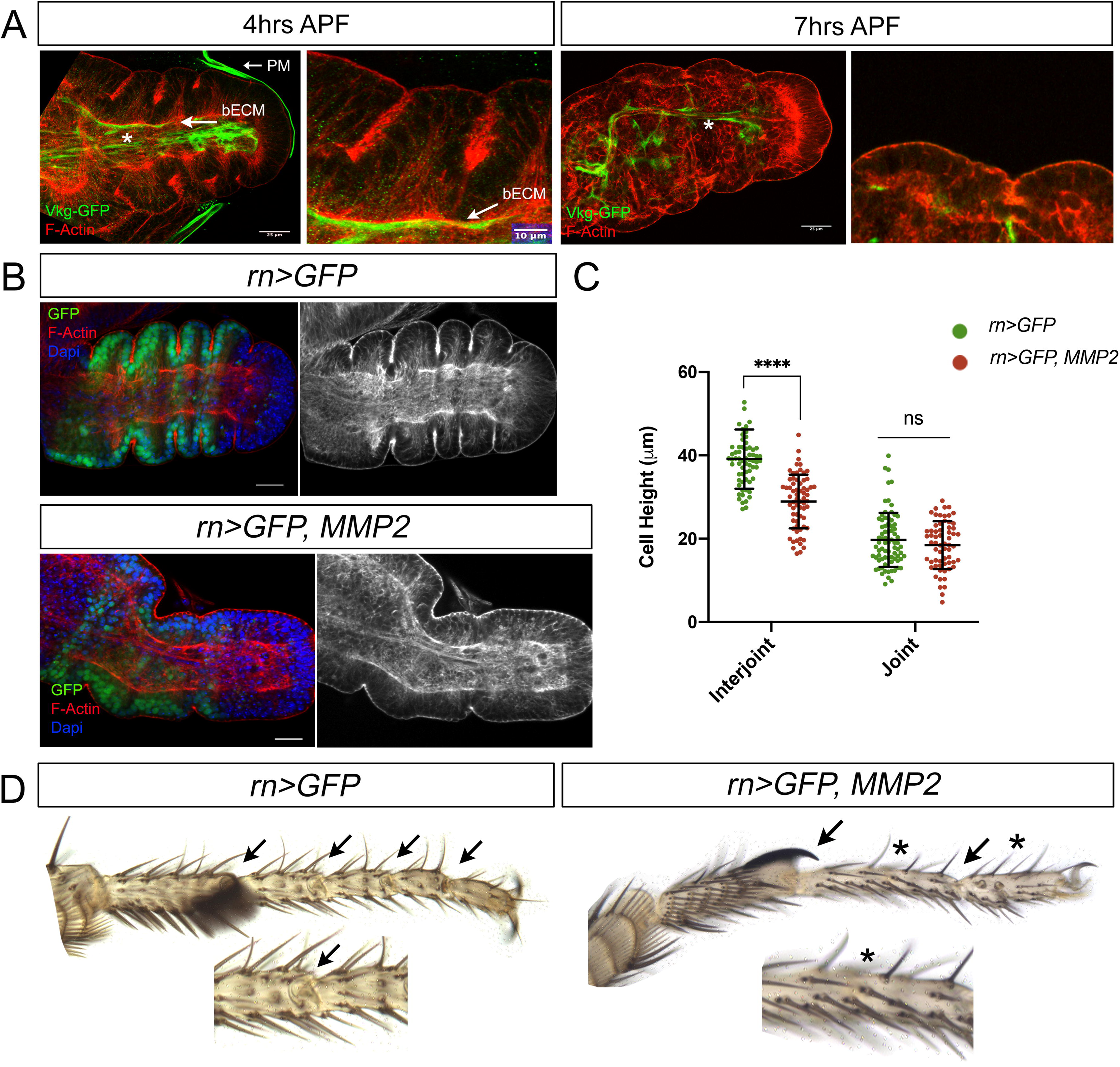
The extracellular matrix contributes to generate the compression forces that promote the folding of the epithelium. A) Tarsal region of prepupal leg imaginal discs dissected 4 hrs APF and 7 hrs APF and stained for F-actin (red) and Vkg-GFP (green). Vkg-GFP covers the basal surface of the main epithelium and of the peripodial membrane at 4 hrs APF. Note the degradation of Vkg-GFP in contact with the epithelial cells and peripodial membrane at 7 hrs APF. * Vkg-GFP also accumulates surrounding the tendon precursors and other internal stuctures. A detailed image of a tarsal joint is shown. The arrow indicates the basal extracellular matrix (bECM) and the peripodial membrane (PM). Scale bar, 25 µm. B) Defective fold formation after the expression of *MMP2* in the tarsal segments of the leg (*rn-Gal4, UAS-GFP*; UAS-*MMP2*). 3-4 hrs prepupae leg discs stained for F-actin (red and gray), GFP (green) and Dapi (blue). Note that the leg is expanded compared to its control (*rn-Gal4, UAS-GFP)*. Scale bar, 25 µm. C) Quantification of the cell heights at the interjoint and joint domains in *rn>GFP* (control, 12 legs, 77 joints, 65 interjoints) and *rn>GFP, MMP2* (12 legs, 63 joints, 66 interjoins) prepupal legs dissected between 3-4 hrs APF and stained for Dysf and F-actin. ****p<0.0001 and ns=not significant, with Student’s t test, indicating a significant difference from control. Error bars represent SD. D) Tarsal segments from adult’s legs expressing the indicated transgenes as in B. Tarsal joints are indicated by arrows and the affected ones by asterisks. A representative close view of a tarsal joint is also presented.

### Dysf induces apical F-actin accumulation and cell shortening

Previous work from our lab have elucidated the role of Dysf, a direct target of Notch, in the formation of both tarsal folds and adult joints (15-17). We have shown that Dysf enhances Rho1 activity, F-actin accumulation and promotes apical constriction at the cells that will form the joint domain (16). This suggests that Dysf confers a unique property to the cells in the joint domain by preventing their apico-basal elongation following tissue compression.

To explore this possibility, we decided to analyze in detail the behavior of the interjoint and joint domain cells after Dysf depletion in the posterior compartment (*hh>GFP, dysf-RNAi)*. As we previously reported, in this experimental condition the posterior compartment loses the characteristic tarsal folds while maintaining Notch activation, visualized by the activity of the Notch target *Enhancer-of-split m*β (*E(spl)m*β) (Fig. 6A) (14, 16, 17). Interestingly, we found that the apico-basal shortening of the corresponding joint cells does not occur in the absence of Dysf (Fig. 6B and C). However, we also noticed that the height of the interjoint cells decreased dramatically when compared to the control anterior domain (Fig. 6B and C).

**Figure 6:**
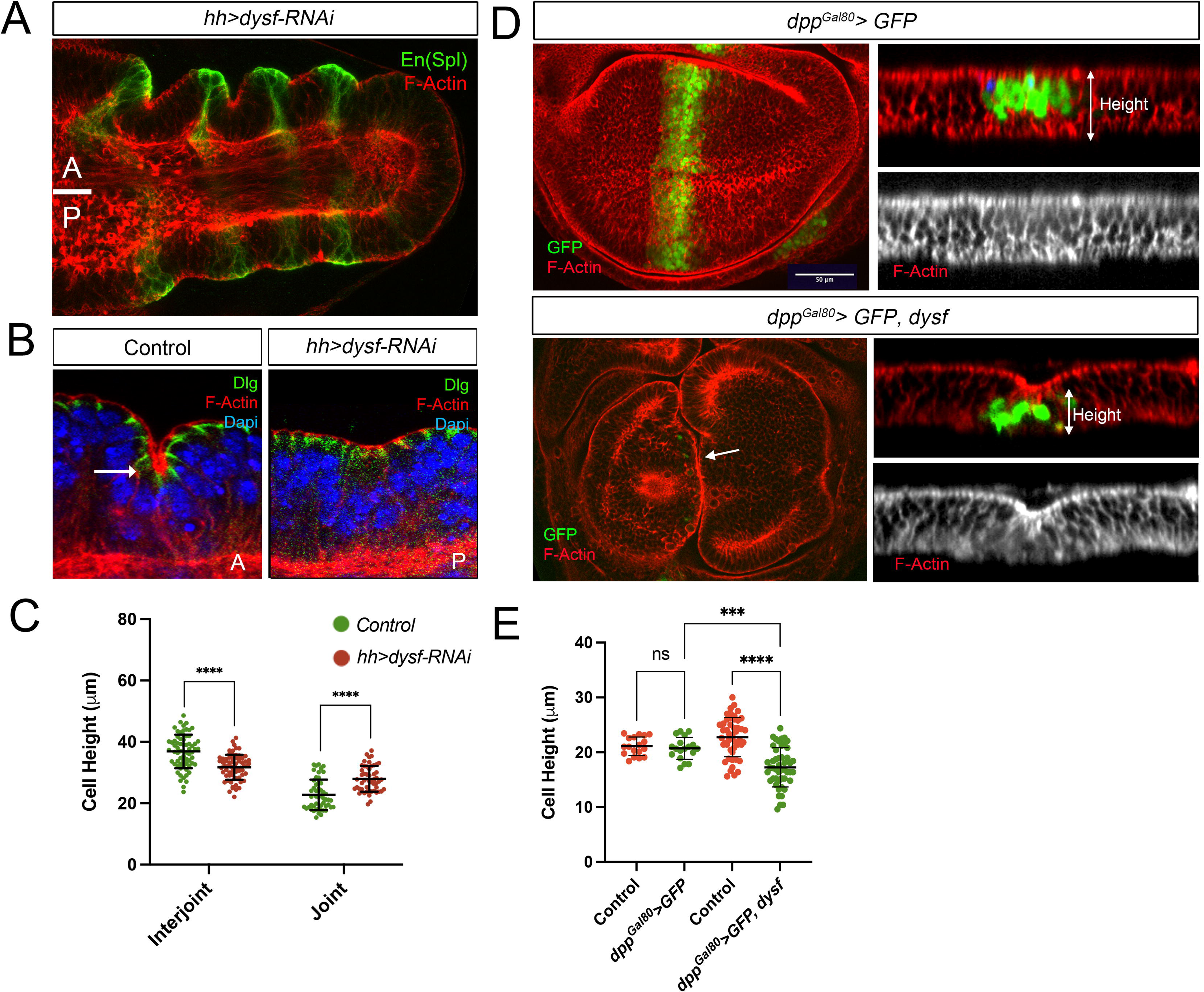
Dysf induces F-actin accumulation and cell shortening. A) Tarsal region of prepupal leg imaginal discs expressing the *UAS-dysf-RNAi* in the posterior compartment by the *hh-Gal4* (*hh>*) stained for F-actin (red) and for the Notch target gene *E(spl)m*β (green). The anterior and posterior compartments are indicated as A and P, respectively and their boundary is represented by white line. B) A higher magnification of an anterior (control) and posterior sagittal view of tarsal folds from a prepupal leg imaginal discs expressing the *UAS-dysf-RNAi* in the posterior compartment. F-actin (red), Dlg (green) and Dapi (blue). An arrow indicates the accumulation of F-actin in the control anterior compartment cells that are apically constricted. C) Quantification of the cell heights at the interjoint and joint domains in *hh>cherry-RNAi* (control, 17 legs, 51 joints, 68 interjoints) and *hh>dysf-RNAi* (17 legs, 51 joints, 68 interjoints) prepupal legs. ****p<0.0001 with Student’s t test, indicating a significant difference from control. Error bars represent SD. D) Apical view and Z-section of a region of the pouch of wing imaginal discs expressing the indicated transgenes for 48 hrs under the *dpp-Gal4, tub-Gal80^ts^* (*dpp^Gal80^>*) line. The *dpp-Gal4* driver was used to express *GFP* and *dysf* in a band of cells of the anterior compartment of the wing pouch. F-actin (red and grays) and GFP (green). The fold generated by the ectopic expression of *dysf* is marked by an arrow. Also indicated is how the cell height is measured. Scale bars: 50 µm. E) Quantification of the cell heights measured in the Z-sections of the genotypes described in D. The region next to the Dpp domain was used as a control for each experiment. *dpp^Gal80^>GFP* (n=31) and *dpp^Gal80^>GFP, dysf* (n=24) Two measurements were done for each condition. Statistical analysis by one-way ANOVA when compared the mean of each genotype with the mean of the control as indicated. ****<p=0.0001, ***<p=0.001 and ns=not significant.

To confirm that Dysf is able to promote the apico-basal shortening of the cells, we ectopically expressed *dysf* with the *dpp-Gal4, tub-Gal80^ts^* (*dpp^Gal80^>*) line in an anterior stripe of cells in the relative flat epithelium of the wing pouch (Fig. 6D). As we previously reported, Dysf is sufficient to induce the apical constriction of the cells, the accumulation of F-actin and the formation of an epithelial fold in the wing pouch (16) (Fig. 6D). Importantly, the cells that ectopically expressed *dysf* have reduced their apico-basal height when compared to their neighbors (Fig. 6D-E).

These and our previous results indicate that Dysf induces Rho1 activity, F-actin accumulation, apical constriction and cell shortening (16).

### Cell proliferation is required for epithelial folding in the wing imaginal disc

As in the leg, the wing imaginal disc is folded in a stereotyped manner. It has been proposed that planar differential rates of cell division initiate the folding of the wing imaginal discs epithelium (4). However, other reports have questioned the relevance of cell proliferation in the formation of the wing disc folds (7). Our results in the leg disc demonstrated that cell division is required for correct epithelial folding and joint formation. Therefore, we tested the consequences of altering cell proliferation for the correct folding of the wing epithelium. First, we knocked down E2f1 or Cdk1 in the posterior compartment with the *hh^Gal80^>* driver during third instar larvae for 48 hrs before dissection and studied the formation of the three main epithelial folds: notum-hinge (NH), hinge-hinge (HH), and hinge-pouch (HP). These folds are readily observed with a F-actin staining in mature third instar larval wing imaginal discs (Figure 7A). In contrast to leg, the wing epithelium is folded at the apical and basal sides. As observed in Fig. 7B and C, altering cell proliferation in the posterior compartment inhibits the correct formation of the HP fold, but not the other two folds (NH and HH). One possibility is that the NH and HH folds do not require cell proliferation for their formation. However, as the HP fold is the last one to be formed (4, 34), it is more likely that this fold is the only one that is affected at the time we reduced proliferation. Second, we tested whether an increase in cell proliferation is sufficient to induce epithelial fold formation. To test this, we temporarily expressed *E2f1+Dp, string* (*stg*)*+CycE* or *yki* for 48 hrs before dissection using the *patched-Gal4, tubGal80ts (ptc^Gal80^>)* driver in the flat epithelium of the wing disc pouch (Fig 7D-G). In order to assess the changes on tissue morphology we stained the discs with Phalloidin. In the control experiment, the expression of GFP in the Ptc domain does not alter wing pouch epithelial morphology (Fig 7D). Overexpression of these factors efficiently induced over proliferation, as visualized by an increase of pH3 mitotic figures with respect to control discs; however, we failed to observe any ectopic fold in the wing pouch apart from a small bulge of the epithelium (Fig 7F-G). These results indicate that although cell proliferation is required for the correct folding of the epithelium, an increase of this parameter is not sufficient to start the process on its own.

**Figure 7:**
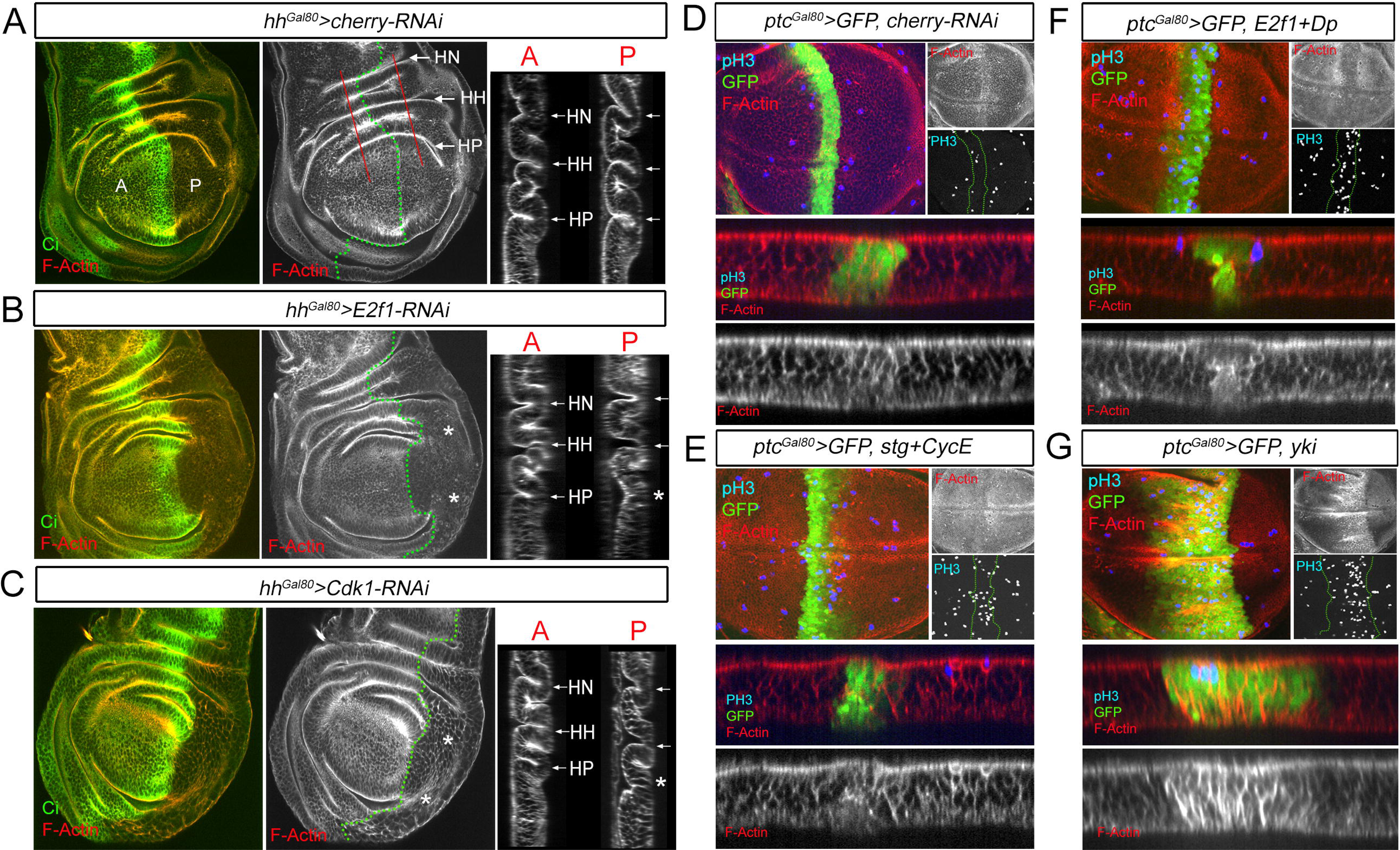
Cell proliferation is required for the correct formation of the hinge-pouch (HP) fold. A-C) Third instar wing imaginal discs stained for F-actin (red and gray) and Ci (green) expressing the indicated transgenes for 36 hrs in the posterior compartment with the *hh-Gal4*, *tub-Gal80^ts^* (*hh^Gal80^>*) driver. The antero-posterior compartment is indicated by a green dotted line and the regions selected for the Z-sections in the anterior (A) and posterior (P) compartment with a red line. The different folds in the wing disc are indicated with arrows as: NH (notum-hinge), HH (hinge-hinge) and HP (hinge-pouch). An asterisk indicates the absence or incomplete formation of a fold. D-G) Apical view of the wing pouch and of a representative Z-section of wing imaginal discs expressing the corresponding transgenes for 48 hrs with the *ptc-Gal4*, *tub-Gal80^ts^* (*hh^Gal80^>*) line. The *ptc-Gal4* driver is used to express different UAS constructs in a band of cells of the anterior compartment of the wing pouch. The imaginal discs were stained for F-actin (red and gray), pH3 (blue and gray) and GFP (green). A dotted green line indicates the Ptc domain.

### Simulation of leg epithelial folding

Our results in the leg discs indicate that cell proliferation, within the environment of the ECM, generates tissue-wide compression forces. This, in collaboration with Dysf activity which directs apical F-actin and cell shortening at the presumptive joint cells, promotes epithelial folding.

To test if the combination of cell proliferation and *dysf* local expression is sufficient to initiate the periodic folding profile that we observe during the early steps of tarsal joint formation, we developed a computer-based simulation model of a simple monolayer of cells set to follow certain basic rules: a) all cells can grow and proliferate; b) dimensions of the system in the x-axis are fixed, so the tissue is not able to expand in this direction. This restriction mimics the effect of the ECM and peripodial membrane; c) each cell grows in volume by compressing its neighbors until they reach a minimal size; d) when neighbors reach this minimal size, cells tend to grow by expanding apically and upwards; e) specific locations in the tissue are defined in such a way that cells inside these domains (marked as red in Fig. 8) cannot grow apically, as occurs with the cells that are influenced by Dysf function in the joint domain. To simplify the model, the red cells (joint) do not reduce their apico-basal length, in contrast to what happens during *in vivo* folding (Fig. 4B and C).

**Figure 8:**
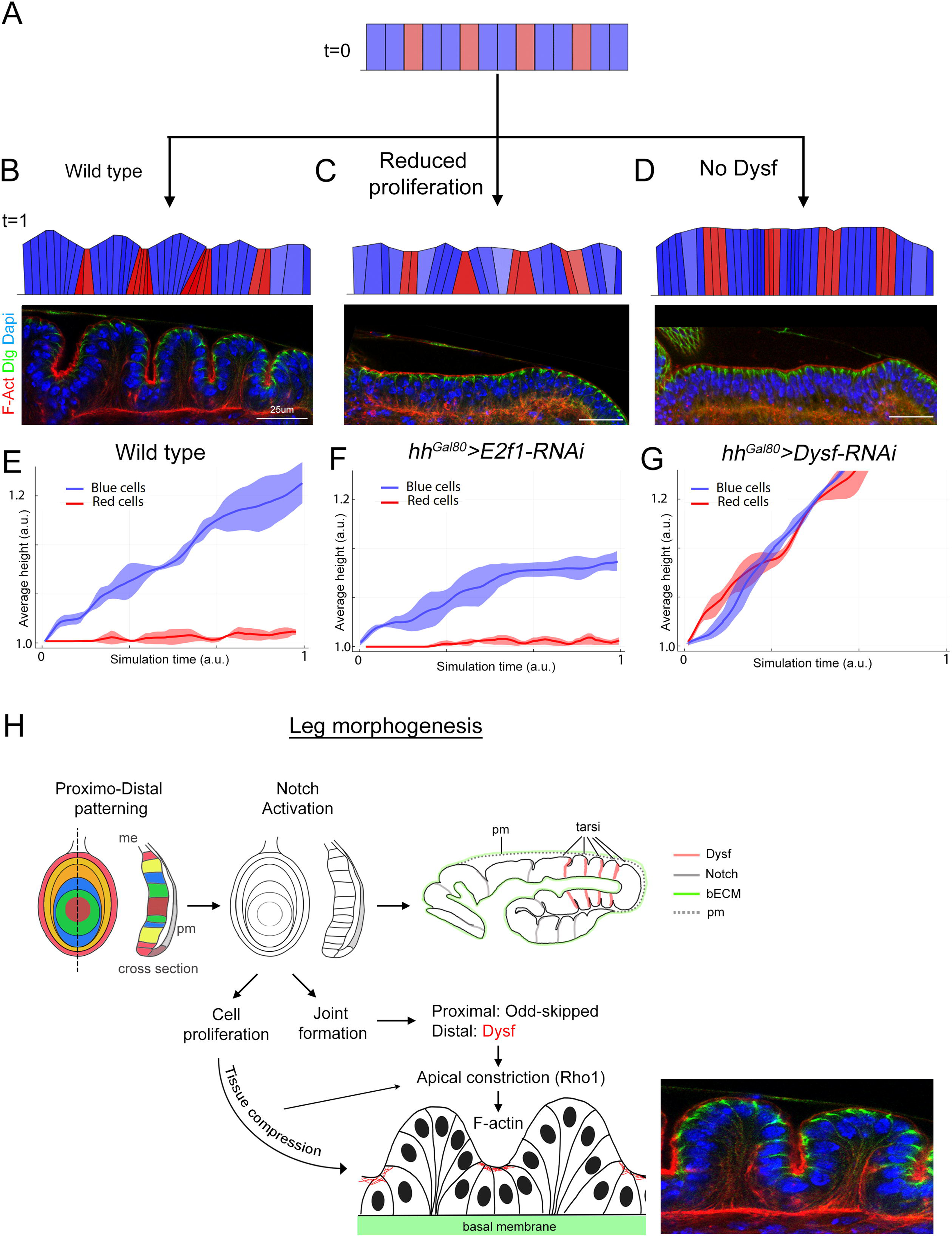
Simulation of leg epithelial folding and proposed model. A) Initial stage (t=0) of the mathematical simulation of the early events of tarsal epithelial folding. The different colors indicate the interjoint (blue) and joint (red) domains. B-D) Final stage of the simulation (t=1) after allowing the cells proliferate normally (B), by reducing proliferation (C) and by removing the restriction to grow apically (D). See the text for full details and Sup. Movies. Under each cartoon the corresponding *in vivo* experiment is presented as a sagittal view of the tarsal epithelial folds that results from knocking down E2f1 or Dysf in the posterior compartment. F-actin (red), Dlg (green) and Dapi (blue). E-G) Quantification of the average blue and red cell heights for each experimental condition of the simulation. The line represents the average of 3 simulations and the ribbon the standard deviation. H) Proposed model for leg epithelial folding. The subdivision of the leg in different domains of gene expression along the PD axis allows the sequential activation of the Notch pathway in concentric rings. Notch is required for the non-autonomous growth of the leg and for the formation of the epithelial folds that prefigure the formation of the adult joints. In the proximal domain, Notch activates the *odd-skypped* family of transcription factors while in the tarsal region induces *dysf* expression. The combined effect of tissue compression generated by the proliferation of leg imaginal cells, within the constraining environment of the ECM and the peripodial membrane, and Dysf promotes the activation of Rho1 and the apical accumulation of F-actin in a specific group of cells, leading to their apical constriction and inability to grow apically (joint cells). In contrast, interjoint cells respond to tissue compression by expanding apically, promoting the buckling of the epithelium. me, main epithelium, pm, peripodial membrane, bECM, basal extracellular matrix. Leg cartoon modified from Kojima, 2017 (50).

With these five very simple assumptions, and starting from a limited number of cells confined in the horizontal direction (Fig. 8A), we run numerical simulations for different conditions (time-lapse movies shown as supplementary material, intensity levels of blue are used to illustrate cell volume). In wild type conditions (Fig. 8B and S1 Movie) cells proliferate increasing the density of the tissue until they reach a maximum when blue cells (interjoint domain cells) start to grow apically compressing the apical domain of red cells (joint domain cells). Since red cells are restricted to grow basally, they expand by compressing the basal domain of their neighbors. The combination of these two processes results in the spontaneous formation of folds in the tissue, that simulate the characteristic folding pattern of the distal leg (Fig. 8B and S1 Movie).

Our experiments provide evidence that a reduction in proliferation or the knockdown of Dysf aborts the formation of folds in the tissue (Fig. 2A and Fig. 6A). To simulate these experimental conditions in our model, we first reduced the average cell cycle length of the cells to levels similar to the knockdown of E2f1 in our experiments (reduction of 35% in the cell number). In these conditions tissue folding is impaired as the cells never reach the confluence required for compressing their neighbors to the extent that they need to expand apically (Fig. 8C and S2 Movie). In this simulation, we observed that joint domain cells (red) suffer less compression that is translated in less apical constriction (S6 Fig). A similar situation is observed in our experiments after E2f1 knockdown, where joint domain cells have lower levels of F-actin accumulation and suffer less apical constriction (Fig. 3B and E).

Second, we model the lack of Dysf by performing simulations where the restriction of apical expansion is eliminated in red cells, and therefore all cells could apically expand if needed as observed in our experiments (Fig. 6A-C). In these conditions, tissue folding is also impaired in the model as no differential apical growth is established between joint and interjoint cells (Fig. 8D and S3 Movie). In addition, we detected the absence of apical constriction in any location of the tissue as observed experimentally (S6 Fig) (16).

## Discussion

In this study, we investigated how cell proliferation, through the accumulation of cell mass, serves as a tissue-wide mechanism, leading to the buckling of the epithelium. During larva development, the leg imaginal disc grows restricted by the ECM and the peripodial membrane, both acting as elastic constraining elements that, together with the accumulation of cells, help generate the compression forces that fold the epithelium at specific positions. In the tarsal segments of the leg these folding positions are defined by the activity of the Notch target gene *dysf*. Importantly, we found that cell proliferation not only contributes to shaping the leg epithelium but is also necessary for the formation of the adult joints.

Previous work by many labs have described precisely the cellular mechanisms that promote the cell shape changes required for epithelial folding (reviewed in (2)). However, the influence that tissue-wide mechanisms have on epithelial folding has been less explored. In the *Drosophila* embryo, a genetic program directs the apical constriction of a group of cells through changes in the acto-myosin network to form the ventral furrow (reviewed in (35)). Importantly, these cell shape changes are coordinated with embryo-wide ectodermal movements towards the furrow that facilitates internalization of mesodermal cells (36, 37). In the wing disc, the position and formation of the epithelial folds is controlled by a patterning genetic network (34, 38-40), that is translated into mechanical forces that changes the shape of the cells that form the folds (7, 41). In addition, differential planar growth rates in the disc in combination with the constraining effects of the basement membrane helps the buckling of the epithelium at precise positions (4). All these studies highlight the importance of tissue-wide forces to act coordinately with local active mechanisms to promote tissue folding and organ shape.

In the developing leg imaginal disc, a combinatorial code of proximo-distal transcription factors locates the expression of the Notch ligands Dl and Ser at the distal end of each leg segment (10-12, 15, 42-44) (Fig. 8H). The activation of Notch not only controls the formation of the adult joints through the regulation of subsidiary transcription factors but is also required for the growth of the leg in a non-autonomous manner (13-15). In the tarsal segments, the Notch target Dysf transcriptionally controls the expression of Rho1 GTPase regulators and of proapoptotic genes at the presumptive tarsal joints (16, 17). The patterned restricted activation of Rho1 promotes actin-myosin network contraction and apical constriction, while localized dying cells at the joint domain generate an apico-basal force that cause the epithelium to fold in a stable manner (5). However, genetic elimination of apoptotic cells has no effect on tarsal epithelial folding nor adult joint formation (16).

Leg imaginal epithelial folds appear in a sequential manner starting at the beginning of the third instar larval period, time of active cell proliferation. Our experiments, which involved reducing cell division rates during the folding process, indicate that a minimal number of cells is required to generate the characteristic folds in the tarsal epithelium. Importantly, the failure to fold the epithelium in these experiments is not due to defects on Notch pathway activation, as the expression of two of its target genes, *dysf* and *bib,* is not affected. Instead, we observed that activation of Rho1 and F-actin accumulation at the joint domain cells, processes that depend on Dysf activity (16), are impaired after reducing cell proliferation rates. These results suggest that tissue compression reinforces Rho1 activation and apical constriction induced by Dysf. Nevertheless, the exact mechanism responsible for this effect remains unknown.

A detailed analysis of the phenotypes resulting from reduced cell numbers uncovers a previously unanticipated behavior of the interjoint cells - their ability to increase in height in response to tissue compression. These data suggest that for the correct formation of the tarsal folds at least two processes act in parallel (Fig. 8H). The first one is an increase of tissue compression due to cell proliferation within the constraining environment of the ECM and the peripodial membrane. Accordingly, basement membrane degradation by the expression of MMP2 in the leg affects fold and joint formation (Fig. 5). The second one is the activation of *dysf* at the distal end of each tarsal segment. Although *dysf* is not expressed in all the cells that form the tarsal fold, its function is absolutely required for the formation of the joint domain (17). We have described that Dysf promotes Rho1 activation and the apical constriction of the cells that is accompanied by a decrease in their height (16). At the same time, cell proliferation generates tissue-wide compression forces that promote different cell behaviors depending on the developmental fate of the cells. The progressive accumulation of cells forces the apico-basal stretch of the interjoint cells, while at the joint domain cells, it facilitates apical constriction through the activation of Rho1 and the apical accumulation of F-actin. Knockdown of E2f1, which leads to a reduction in cell number, suppresses the increase in cell height of interjoint cells and reduces the apical constriction of joint cells. This results in the elimination of the fold and the adult joint. Other mechanisms besides cell proliferation could also increase tissue compression and contribute to the formation of the epithelial folds such as oriented cell division, cell intercalation and cell shape changes (22, 45, 46).

Using the wing disc, we were also able to demonstrate that cell proliferation is required, at least, for the formation of the HP fold. Interestingly, non-uniform growth rates contribute to the correct location of the folds at the time of their formation, confirming that cell proliferation and the genetic patterning mechanisms act coordinately to shape a tissue (4).

In this work we also recreated the initial steps of tarsal fold formation by a simple computer-based simulation model, in which the pattern of Dysf function is generated by preventing a group of cells to grow apically due to F-actin accumulation. In this model, all cells proliferate in a confined space, as it happens *in vivo* by the presence of the ECM and the peripodial membrane. The cells compress their neighbors until they reach a minimum volume, ultimately resulting in the formation of folds. As it occurs *in vivo*, interjoint cells start to increase in height while joint cells progressively constrict their apical side and expand basally generating the folding observed in the tarsal segments. Remarkably, this simple model predicts the folding phenotypes and the behavior of the cells after reducing cell proliferation and in the absence of *dysf*.

In summary, this work emphasizes the importance of studying both local and tissue-wide mechanisms that together orchestrate the formation of epithelial folds and organ architecture in a reproducible pattern.

## Materials and methods

### *Drosophila* melanogaster lines

The culture of *Drosophila* melanogaster strains was done in light/dark cycles of 12 hrs in incubation chambers under controlled temperature (17°C, 25°C or 31°C, depending on the experiments).

The following Gal4 lines were used and described in Flybase except when noted: *ptc-Gal4*, *dpp-Gal4*, *rn-Gal4*, *Dll^212^-Gal4*, *ap^42B11^-Gal4 and dysf-Gal4* (Flylight database: http://flweb.janelia.org/cgi-bin/flew. cgi) (47), *hh-Gal4*, *bib-Gal4*. To temporarily restrict the activity of the different Gal4 lines we used the tubulin-Gal80ts system. Briefly, embryos were collected for 24 to 48 hrs, maintained at the restrictive temperature (17° C) and then shifted to the permissive temperature (31° C) for the required time prior to dissection.

The following UAS lines were used: *UAS-GFP*, *UAS-cherry-RNAi* (used as control, BDSC#35785), *UAS-MMP2* (BDSC#58705), *UAS-dysf-RNAi* (VDRC#110381), *UAS-CycE-RNAi* (VDRC#110204), *UAS-E2f1-RNAi* (VDRC#108837), *UAS-Cdk1-RNAi* (VDRC#106130 and BDSC#28368)*, UAS-E2f1+Dp* (BDSC#4774 and #4770)*, UAS-miRHG* (29)*, UAS-yki* (48)*, UAS-Rbf^280^*(BDSC#50748), UAS-*Rho1-BD-GFP*(27), *UAS-dysf* (49), *UAS-stg* (BDSC#4777) and *UAS-CycE* (BDSC#4781).

Some of the RNAi lines were combined with *UAS-Dcr2* (BDSC#24646) to enhance its effect.

Other reporter and GFP protein trapped lines used are: *E-cad-GFP* (BDSC#60584), *vkg-GFP* (FlyTrap G205), *bib-lacZ* and *E(spl)-m*β*-CD2* (14), *dysf-lacZ* (17), *E2f1-GFP* (BDSC#83388) and *Sqh-GFP* (gift from M. Suzanne).

### Immunostaining

Standard procedures were used to fix and stain prepupal (from 0 to 12 hrs APF) and larval leg and wing imaginal discs. Briefly, larvae and prepupae were dissected in PBS and fixed with 4% paraformaldehyde in PBS for 25 minutes at room temperature. They were blocked in PBS, 1% BSA, 0.3% Triton for 1 hr, incubated with the primary antibody over night at 4 C, washed four times in blocking buffer, and incubated with the appropriate fluorescent secondary antibodies for 1.5 hours at room temperature in the dark. They were then washed and mounted in Vectashield (Cat#H-1000). We used anti-Phalloidin (TRITC) (Sigma-Aldrich Cat#P1951) to stain the actin cytoskeleton, and TOPRO (Thermo-Fisher Cat#T3605) or Dapi (MERCK) to stain nuclei.

As primary antibodies, we used mouse anti-Dlg (DSHB Cat# 4F3, 1:50), rat anti-Ser (a gift from Ken Irvine, Rutgers University, 1/1000), mouse anti-En (DSHB Cat# 4D9, 1:50), rabbit and mouse anti-β-Gal (MP Biomedics #559761 and Promega #Z378A, 1:1000), rabbit and mouse anti-pH3 (Merck Millipore #06-570 and Cell signal technology #9796, 1/500) and rat anti-Ci (DSHB 2A1, 1/50).

To preserve the 3D structure of the imaginal discs, the samples were mounted using an adhesive spacer at both sides of the samples in the slide. A coverslip was then placed on top of the spacer.

When indicated, prepupae were synchronized to properly compare fold formation phenotypes. White pupae were selected of the given phenotype, incubated for 0, 3, 5 and 6 hrs at the required temperature and then dissected and stained following standard procedures.

For Edu staining, dissected leg discs were cultured in 1 mL of EdU labelling solution for 20 min at room temperature and subsequently fixed in 4% paraformaldehyde for 30 min at room temperature. EdU detection was performed according to the manufacturer instructions (Click-iT EdU Alexa Fluor Imaging Kit, ThermoFisher Scientific), and leg discs were incubated during 30-40 min at room temperature in the dark.

All confocal images were obtained using a Leica LSM710 vertical confocal microscope and were treated using Fiji and Photoshop programs.

Rho1RBD-GFP relative fluorescence intensity was quantified by measuring the mean intensity of fold and interjoint domain cells and their areas were calculated manually using ROI measurement tool in Fiji. Apical F-actin fluorescence intensity was measured in the joint domain cells or in the corresponding domain in the experimental compartment using Fiji. Sagittal images were obtained for prepupal leg imaginal discs. Cross sectional images from wing imaginal discs were obtained by Fiji from multiple Z-stacks that cover the whole disc.

### Adult leg preparations

Adult or pharate (in the case of flies that could not hatch) legs of the required phenotypes were collected in 96% ethanol until mounted. We used Hoyer’s mounting medium in a1:1 proportion with lactic acid (90% MERCK) to preserve the cuticle of the legs. Multiple focal planes of each leg were acquired and then combined using the Helicon Focus program to create a fully focused image of the legs.

### Statistical analysis

Statistical analysis was performed using the GraphPad Prism software (https://www.graphpad.com). To compare between two groups, a non-parametric Student’s *t*-test test was used. To compare between more than two groups, a non-parametric, one-way ANOVA Dunnett’s test was used. Sample size was indicated in each figure legend.

### Simulation of leg epithelial folding

For the simulations, we developed a simplified agent-base model written in Julia language as a Jupyter notebook (available as supplementary material), using the Plots (for illustrations) and RollingFunctions (for smoothing and average functions). In the model, cells are numerical entities with spatial dimensions defined by the user, organized as a one-dimensional array that simulates a monolayer, embedded in a domain of fixed length. The model is a time-based algorithm with a for cycle that sets the total number of iterations and the time-step. For each iteration, a single cell in the tissue is chosen at random, that is set to grow and/or proliferate based on the following rules:

- Cells grow by expanding any of the four vertices. The direction of growth is chosen at random. Since the domain size is fixed, cells grow by compressing their neighboring cells: Therefore, to maintain the integrity of the tissue, when a given cell is set to grow in a given direction, the adjacent cell is compressed accordingly.
- A minimal cell size is defined in such a way that a cell cannot be compress below this value. In consequence, a cell selected to grow in the close vicinity has to expand in other way.
- Two types of cells are defined: cells that can grow both horizontally and vertically (labeled in blue) and cells that, due to elevated levels of actin, cannot grow vertically (labeled in red). Color intensity of the is set to illustrate the compression level of each cell (light color correspond to large cells, and dark color corresponds to a cell highly compressed close to its minimal size). Therefore, red cells close to a highly compressed cell tend to expand apically, while blue cells set to grow against a highly compressed neighbor expand basally.
- The cell cycle length (defined as the number of iterations for a cell to enter mitosis and divide) is set by the user. When a given cell is older than this value, it divides in half producing to new cells of the same type, that occupy the same space as their mother.

The state of the system in terms of size and position of each cell is recorded at each time step and draw to generate the time-lapse movies for each condition.

Code is available as a Jupyter notebook in supplemental protocol.

## Acknowledgements

The authors would like to thank the Bloomington Stock Center, the Vienna Drosophila Resource Center, and the Developmental Studies Hybridoma Bank for fly stocks and reagents. The authors specially thank the Confocal Microscopy Service at CBMSO and Ernesto Sanchez-Herrero, Jose Felix de Celis and the people from the lab for fruitful discussions throughout the period of this work.

This study was supported by grants from: FEDER/Ministerio de Ciencia e Innovación-Agencia Estatal de Investigación-Consejo Superior de Investigaciones Científicas (No. PGC2018-095144-B-I00, PID2021-127114NB-100 to CE).

---

**S1 Fig: Cell proliferation and *dysf* expression during leg development.**

A and B) Third instar (A) and a distal prepupal (B) leg imaginal disc stained for EdU (red and gray) and *dysf-lacZ* (green). ta, tarsal segment.

**S2 Fig: Downregulation of E2f1 in leg and wing imaginal discs.**

A) *E2f1-GFP* (green) expression in third instar leg and wing imaginal discs.

B) Leg and wing imaginal discs of the *hh^Gal80^>E2f1-RNAi* genotype dissected 48 hrs after inducing the transgene in the posterior compartment and stained for E2f1-GFP (green) and Ci (red).

C) Wing imaginal discs expressing the indicated transgene with the *hh^Gal80^>* dissected 48 hrs after inducing the transgene in the posterior compartment and stained for Ci (green) and pH3 (red).

D) Quantification of the size effect phenotype produced by the downregulation of E2f1 in the posterior compartment as in C presented as the ratio between the posterior and anterior area. ***p<0.001 with Student’s t test, indicating a significant difference from control. Error bars represent the standard error of the mean (SEM).

E) Mitotic index measured as the number of pH3-positive cells per area in the posterior domain of control (*hh^Gal80^>cherry-RNAi*) and E2f1 knockdown (*hh^Gal80^>cherry-RNAi*). ns=not significant with Student’s t test, indicating a not significant difference from control. Error bars represent the standard error of the mean (SEM).

**S3 Fig: Fold and adult joint phenotypes after reducing cell proliferation in the leg.**

A) Third instar leg and prepupal leg imaginal discs stained for Dlg (red) or F-actin (red) and the *ap^42B11^>GFP* line (green). A higher magnification of a sagittal view of the last tarsal segments showing the ap domain that encompasses the fold between the fourth and fifth tarsal segments (GFP, green).

B) Adult legs expressing the indicated transgenes with the *ap^42B11^*> driver. The tarsal segments and a higher magnification of the last segments is shown. An arrow point to the ta4/ta5 joint and an asterisk indicates the absence of the joint.

C) Distal region of 3-4 hrs APF leg discs stained for F-actin (red) and *dysf-lacZ* (green) expressing the indicated transgenes with the *ap^42B11^*> driver. The fold between the fourth and fifth tarsal segments is indicated with an arrow and its absence or defective formation with an asterisk.

D) Quantification of the ta4/ta5 adult joint phenotypes for the genotypes indicated. The number of adult legs scored are: *ap^42B11^>cherry-RNAi*: 48, *ap^42B11^>E2f1-RNAi*: 65, *ap^42B11^>Rbf^280^*: 53 and *ap^42B11^>CycE-RNAi*: 57.

**S4 Fig: Knocking down E2f1 in the joint domain.**

A) Tarsal region of prepupal leg imaginal discs expressing the indicated transgenes with the *dysf-Gal4* line stained for F-actin (red and gray) and for GFP (green).

B) Tarsal segments from adult’s legs of the genotypes in A, expressing the corresponding transgenes.

C) Tarsal segments from adult’s legs expressing the indicated transgenes with the bib-Gal4 line.

**S5 Fig: Temporal expression of *yki* in the posterior compartment.**

A) Prepupal leg imaginal discs of the *hh^Gal80^>yki* genotype dissected 24-30 hrs after inducing the transgene in the posterior compartment and stained for F-actin (red and gray) and Ci (green). The antero-posterior compartment boundary is represented by a white line. A higher magnification of anterior (A) and posterior (P) tarsal folds is indicated. The anterior compartment is used as a control. Also indicated is how the joints depth is measured.

B) Quantification of joint depth and of the cell heights at the interjoint and joint domains of the prepupal legs in A. A total of 19 legs were dissected and 56 joints and 75 interjoints were measured. ****p<0.0001 and *p<0.05 with Student’s t test, indicating a significant difference from control. Error bars represent the minimum and maximum points.

**S6 Fig: Simulation of leg epithelial folding and apical width measurements.**

A) Initial stage (t=0) of the mathematical simulation of the early events of tarsal epithelial folding. The different colors indicate the interjoint (blue) and joint (red) domains.

B-D) Final stage of the simulation (t=1) after allowing the cells proliferate normally (B), by reducing proliferation (C) and by removing the restriction to grow apically (D). See the text for full details.

E-G) Quantification of the average apical width of the red cells for each experimental condition of the simulation.

**S1 Movie:**

Giff movie representing the computer-based simulation of the early events of tarsal epithelial folding. The different colors indicate the interjoint (blue) and joint (red) domains.

**S2 Movie:**

Giff movie representing the computer-based simulation of the early events of tarsal epithelial folding after reducing cell proliferation. The different colors indicate the interjoint (blue) and joint (red) domains.

**S3 Movie:**

Giff movie representing the computer-based simulation of the early events of tarsal epithelial folding after removing the restriction to grow apically as it happens in a *dysf* mutant. The different colors indicate the interjoint (blue) and joint (red) domains.

**S1 Protocol:** Code for the computer-based simulation model model written in Julia language as a Jupyter notebook.

## References

1. Zartman JJ, Shvartsman SY. Unit operations of tissue development: epithelial folding. Annu Rev Chem Biomol Eng. 2010;1:231–46.

2. Tozluoglu M, Mao Y. On folding morphogenesis, a mechanical problem. Philos Trans R Soc Lond B Biol Sci. 2020;375(1809):20190564.

3. Martin AC, Goldstein B. Apical constriction: themes and variations on a cellular mechanism driving morphogenesis. Development. 2014;141(10):1987–98.

4. Tozluoglu M, Duda M, Kirkland NJ, Barrientos R, Burden JJ, Munoz JJ, et al. Planar Differential Growth Rates Initiate Precise Fold Positions in Complex Epithelia. Developmental cell. 2019;51(3):299–312 e4.

5. Monier B, Gettings M, Gay G, Mangeat T, Schott S, Guarner A, et al. Apico-basal forces exerted by apoptotic cells drive epithelium folding. Nature. 2015;518(7538):245-8.

6. Kondo T, Hayashi S. Mitotic cell rounding accelerates epithelial invagination. Nature. 2013;494(7435):125-9.

7. Sui L, Alt S, Weigert M, Dye N, Eaton S, Jug F, et al. Differential lateral and basal tension drive folding of Drosophila wing discs through two distinct mechanisms. Nat Commun. 2018;9(1):4620.

8. Trushko A, Di Meglio I, Merzouki A, Blanch-Mercader C, Abuhattum S, Guck J, et al. Buckling of an Epithelium Growing under Spherical Confinement. Developmental cell. 2020;54(5):655–68 e6.

9. Savin T, Kurpios NA, Shyer AE, Florescu P, Liang H, Mahadevan L, et al. On the growth and form of the gut. Nature. 2011;476(7358):57-62.

10. Ruiz-Losada M, Blom-Dahl D, Cordoba S, Estella C. Specification and Patterning of Drosophila Appendages. J Dev Biol. 2018;6(3).

11. Estella C, Voutev R, Mann RS. A dynamic network of morphogens and transcription factors patterns the fly leg. Current topics in developmental biology. 2012;98:173–98.

12. Rauskolb C. The establishment of segmentation in the Drosophila leg. Development. 2001;128(22):4511–21.

13. Rauskolb C, Irvine KD. Notch-mediated segmentation and growth control of the Drosophila leg. Developmental biology. 1999;210(2):339–50.

14. de Celis JF, Tyler DM, de Celis J, Bray SJ. Notch signalling mediates segmentation of the Drosophila leg. Development. 1998;125(23):4617–26.

15. Cordoba S, Estella C. Role of Notch Signaling in Leg Development in Drosophila melanogaster. Adv Exp Med Biol. 2020;1218:103–27.

16. Cordoba S, Estella C. The transcription factor Dysfusion promotes fold and joint morphogenesis through regulation of Rho1. PLoS genetics. 2018;14(8):e1007584.

17. Cordoba S, Estella C. The bHLH-PAS transcription factor dysfusion regulates tarsal joint formation in response to Notch activity during drosophila leg development. PLoS genetics. 2014;10(10):e1004621.

18. Monier B, Suzanne M. The Morphogenetic Role of Apoptosis. Current topics in developmental biology. 2015;114:335–62.

19. Manjon C, Sanchez-Herrero E, Suzanne M. Sharp boundaries of Dpp signalling trigger local cell death required for Drosophila leg morphogenesis. Nat Cell Biol. 2007;9(1):57–63.

20. Diaz-de-la-Loza MD, Ray RP, Ganguly PS, Alt S, Davis JR, Hoppe A, et al. Apical and Basal Matrix Remodeling Control Epithelial Morphogenesis. Developmental cell. 2018;46(1):23–39 e5.

21. Pastor-Pareja JC, Xu T. Shaping cells and organs in Drosophila by opposing roles of fat body-secreted Collagen IV and perlecan. Developmental cell. 2011;21(2):245–56.

22. Condic ML, Fristrom D, Fristrom JW. Apical cell shape changes during Drosophila imaginal leg disc elongation: a novel morphogenetic mechanism. Development. 1991;111(1):23–33.

23. Ohtani K, DeGregori J, Nevins JR. Regulation of the cyclin E gene by transcription factor E2F1. Proceedings of the National Academy of Sciences of the United States of America. 1995;92(26):12146–50.

24. Xin S, Weng L, Xu J, Du W. The role of RBF in developmentally regulated cell proliferation in the eye disc and in Cyclin D/Cdk4 induced cellular growth. Development. 2002;129(6):1345–56.

25. Gho M, Bellaiche Y, Schweisguth F. Revisiting the Drosophila microchaete lineage: a novel intrinsically asymmetric cell division generates a glial cell. Development. 1999;126(16):3573–84.

26. Tajiri R, Misaki K, Yonemura S, Hayashi S. Joint morphology in the insect leg: evolutionary history inferred from Notch loss-of-function phenotypes in Drosophila. Development. 2011;138(21):4621–6.

27. Simoes S, Denholm B, Azevedo D, Sotillos S, Martin P, Skaer H, et al. Compartmentalisation of Rho regulators directs cell invagination during tissue morphogenesis. Development. 2006;133(21):4257–67.

28. Proag A, Monier B, Suzanne M. Physical and functional cell-matrix uncoupling in a developing tissue under tension. Development. 2019;146(11).

29. Siegrist SE, Haque NS, Chen CH, Hay BA, Hariharan IK. Inactivation of both Foxo and reaper promotes long-term adult neurogenesis in Drosophila. Current biology: CB. 2010;20(7):643–8.

30. Moon NS, Di Stefano L, Morris EJ, Patel R, White K, Dyson NJ. E2F and p53 induce apoptosis independently during Drosophila development but intersect in the context of DNA damage. PLoS genetics. 2008;4(8):e1000153.

31. Huang J, Wu S, Barrera J, Matthews K, Pan D. The Hippo signaling pathway coordinately regulates cell proliferation and apoptosis by inactivating Yorkie, the Drosophila Homolog of YAP. Cell. 2005;122(3):421–34.

32. Harmansa S, Erlich A, Eloy C, Zurlo G, Lecuit T. Growth anisotropy of the extracellular matrix shapes a developing organ. Nat Commun. 2023;14(1):1220.

33. Milner MJ, Bleasby AJ, Kelly SL. The role of the peripodial membrane of leg and wing imaginal discs ofDrosophila melanogaster during evagination and differentiation in vitro. Wilehm Roux Arch Dev Biol. 1984;193(3):180–6.

34. Sui L, Pflugfelder GO, Shen J. The Dorsocross T-box transcription factors promote tissue morphogenesis in the Drosophila wing imaginal disc. Development. 2012;139(15):2773–82.

35. Gheisari E, Aakhte M, Muller HJ. Gastrulation in Drosophila melanogaster: Genetic control, cellular basis and biomechanics. Mechanisms of development. 2020;163:103629.

36. Rauzi M, Krzic U, Saunders TE, Krajnc M, Ziherl P, Hufnagel L, et al. Embryo-scale tissue mechanics during Drosophila gastrulation movements. Nat Commun. 2015;6:8677.

37. Guo H, Huang S, He B. Evidence for a Role of the Lateral Ectoderm in Drosophila Mesoderm Invagination. Front Cell Dev Biol. 2022;10:867438.

38. Sui L, Dahmann C. Wingless counteracts epithelial folding by increasing mechanical tension at basal cell edges in Drosophila. Development. 2020;147(5).

39. Villa-Cuesta E, Gonzalez-Perez E, Modolell J. Apposition of iroquois expressing and non-expressing cells leads to cell sorting and fold formation in the Drosophila imaginal wing disc. BMC Dev Biol. 2007;7:106.

40. Wang D, Li L, Lu J, Liu S, Shen J. Complementary expression of optomotor-blind and the Iroquois complex promotes fold formation to separate wing notum and hinge territories. Developmental biology. 2016;416(1):225–34.

41. Sui L, Dahmann C. Increased lateral tension is sufficient for epithelial folding in Drosophila. Development. 2020;147(23).

42. Estella C, Mann RS. Logic of Wg and Dpp induction of distal and medial fates in the Drosophila leg. Development. 2008;135(4):627–36.

43. Galindo MI, Bishop SA, Greig S, Couso JP. Leg patterning driven by proximal-distal interactions and EGFR signaling. Science. 2002;297(5579):256-9.

44. Campbell G. Distalization of the Drosophila leg by graded EGF-receptor activity. Nature. 2002;418(15):781–5.

45. Taylor J, Adler PN. Cell rearrangement and cell division during the tissue level morphogenesis of evaginating Drosophila imaginal discs. Developmental biology. 2008;313(2):739–51.

46. Baena-Lopez LA, Baonza A, Garcia-Bellido A. The orientation of cell divisions determines the shape of Drosophila organs. Current biology: CB. 2005;15(18):1640–4.

47. Jory A, Estella C, Giorgianni MW, Slattery M, Laverty TR, Rubin GM, et al. A survey of 6,300 genomic fragments for cis-regulatory activity in the imaginal discs of Drosophila melanogaster. Cell reports. 2012;2(4):1014–24.

48. Zhang L, Ren F, Zhang Q, Chen Y, Wang B, Jiang J. The TEAD/TEF family of transcription factor Scalloped mediates Hippo signaling in organ size control. Developmental cell. 2008;14(3):377–87.

49. Jiang L, Crews ST. The Drosophila dysfusion basic helix-loop-helix (bHLH)-PAS gene controls tracheal fusion and levels of the trachealess bHLH-PAS protein. Molecular and cellular biology. 2003;23(16):5625–37.

50. Kojima T. Developmental mechanism of the tarsus in insect legs. Curr Opin Insect Sci. 2017;19:36–42.

